# Violated predictions enhance the representational fidelity of visual features in perception

**DOI:** 10.1101/2024.03.27.587109

**Authors:** Reuben Rideaux, Phuong Dang, Luke Jackel-David, Zak Buhmann, Dragan Rangelov, Jason B Mattingley

**Affiliations:** School of Psychology, The University of Sydney, Camperdown, Australia; Queensland Brain Institute, The University of Queensland, St Lucia, Australia; School of Psychology, The University of Queensland, St Lucia, Australia; Department of Psychological Sciences, Swinburne University of Technology, Hawthorn, Australia

## Abstract

Predictive coding theories argue that recent experience establishes expectations that generate prediction errors when violated. In humans, brain imaging studies have revealed unique signatures of violated predictions in sensory cortex, but the perceptual consequences of these effects remain unknown. We had observers perform a dual-report task on the orientation of a briefly presented target grating within predictable or random sequences, while we recorded pupil size as an index of surprise. Observers first made a speeded response to categorize the orientation of the target grating (clockwise or counterclockwise from vertical), then reproduced its orientation without time pressure by rotating a bar. This allowed us to separately assess response speed and precision for the same stimuli. Critically, on half the trials, the target orientation deviated from the spatiotemporal structure established by the preceding gratings. Observers responded faster and more accurately to unexpected gratings, and pupillometry provided physiological evidence of observers’ surprise in response to these events. In a second experiment, we cued the spatial location and timing of the grating and found the same pattern of results, demonstrating that unexpected orientation information is sufficient to produce faster and more precise responses, even when the location and timing of the relevant stimuli are fully expected. These findings indicate that unexpected events are prioritized by the visual system both in terms of processing speed and representational fidelity.

## INTRODUCTION

Predictive coding theories argue that *prediction errors* are generated when bottom-up sensory inputs deviate from top-down expectations (Friston, 2005). There is broad consensus that prediction errors are associated with increased neural activation (Alink et al., 2010; Den Ouden et al., 2012; Kok et al., 2012; Meyer & Olson, 2011; Richter et al., 2018; Todorovic et al., 2011), which is typically observed during the initial processing cascade in sensory cortices associated with the unexpected stimulus (Tang et al., 2018, 2023), but has also been reported in subcortical regions (Mazancieux et al., 2023). By contrast, it remains unclear how these modulations reshape the pattern activity that represents the unexpected stimulus.

One might hypothesize that expected events should produce more precise representations, because the brain can anticipate incoming sensory signals and provide an optimized neural environment to receive them; for example, by increasing the gain of neurons tuned to the expected stimulus features. Such benefits of anticipation are evident in spatiotemporal cueing (Rohenkohl et al., 2014), and in sequential dependency experiments, where observers respond faster and more accurately to events that comply with a recent history of alternations and repetitions between two states (e.g., reporting on state “A” after exposure to states ABAB) than to those that violate this pattern (e.g., reporting on state “A” after exposure to states BABA; Remington, 1969). There is also behavioural evidence that memory recall is more precise for expected visual objects then unexpected ones (Abreo et al., 2023), and that expected stimuli are more likely to be consciously perceived than competing unexpected stimuli during binocular rivalry (Denison et al., 2011; Pinto et al., 2015).

By contrast, it seems equally reasonable to hypothesize that *unexpected* events might produce more precise representations. Unexpected or ‘surprising’ events are more likely to have harmful consequences than expected ones precisely because they have not been anticipated and thus an appropriate response has not been prepared. On this account, prioritizing the speed and fidelity^1^ with which unexpected events are represented may have adaptive value. Similarly, prioritizing unexpected events to effectively update internal predictive models may support adaptive future behaviours, e.g., increased caution after narrowly avoiding an obstruction on the road (Friston, 2009; Soltani & Izquierdo, 2019). Some behavioural evidence for prioritization of unexpected events has been reported in studies of representational momentum (Hubbard, 1994, 2005) and the von Restorff effect (Wallace, 1965), in which incongruent or “distinct” visual stimuli benefit from improved recall.

Neuroimaging studies addressing the question of how prediction errors shape sensory representations have produced inconsistent findings. Some evidence from functional magnetic resonance imaging (fMRI) suggests that neural representations formed by unexpected stimuli are less precise than those of expected stimuli (Kok et al., 2012), while others suggest the opposite (Richter et al., 2022). Evidence from electroencephalography (EEG) in humans and two-photon calcium (Ca^2+^) imaging in mice indicates that neural representations of unexpected stimuli are more precise than those of expected stimuli (Tang et al., 2018, 2023), while magnetencephalography work has suggested the opposite (Kok et al., 2017). Adding further uncertainty to this question, a recent study failed to find a difference between the neural representations formed by expected and unexpected stimuli (den Ouden et al., 2023).

The hierarchical nature of predictive coding, combined with the multitude of approaches for studying brain function, likely contributes to the inconsistent findings on how prediction errors shape sensory representations. These discrepancies might be due to stimulus and experimental design choices (e.g., some designs may not sufficiently establish sensory predictions), but they might also be explained by biophysical differences between neural recording techniques (e.g., fMRI and EEG) or analytic tools for estimating representational precision from neural activity (e.g., linear discriminant analysis versus inverted encoding) (Brouwer & Heeger, 2009; Harrison et al., 2023; Rideaux et al., 2023).

To test the influence of prediction errors on the representation of visual stimuli, here we sought to measure perception directly, rather than indirectly via neural recordings. In Experiment 1, we had observers (N = 60 adults) perform a dual-report task on the orientation of a briefly presented target grating embedded within sequences that were either predictable (consistent clockwise or counterclockwise rotation) or random (see **Fig. 1**). A similar presentation sequence was used previously to measure prediction errors in single neurons in mouse visual cortex (Tang et al., 2023). Observers first made a speeded binary response to categorize the orientation of the target grating (clockwise or counterclockwise from vertical), then reproduced its orientation without time pressure by rotating a bar. This allowed us to separately assess response speed and precision for the same stimulus. Critically, while the target grating orientation was selected at random, the orientations of the preceding gratings were either sequential (rotating at 30° per grating) or random. Thus, in the sequential condition, an expectation was established and subsequently violated by the target grating. Note, although the target grating in the sequential condition violated the sequence on most trials, this remained an unexpected event relative to the multiple preceding gratings in the trial that conformed to the sequence. By contrast, in the random condition observers had no stimulus information on which to establish an expectation of grating orientation (**Fig. 1a**). To isolate the contribution of the feature-specific violation, we ran a further study (Experiment 2) in which the gratings not only rotated but also traversed an imaginary circle, such that the target grating was either presented at a cued or uncued location.

**Figure 1.**
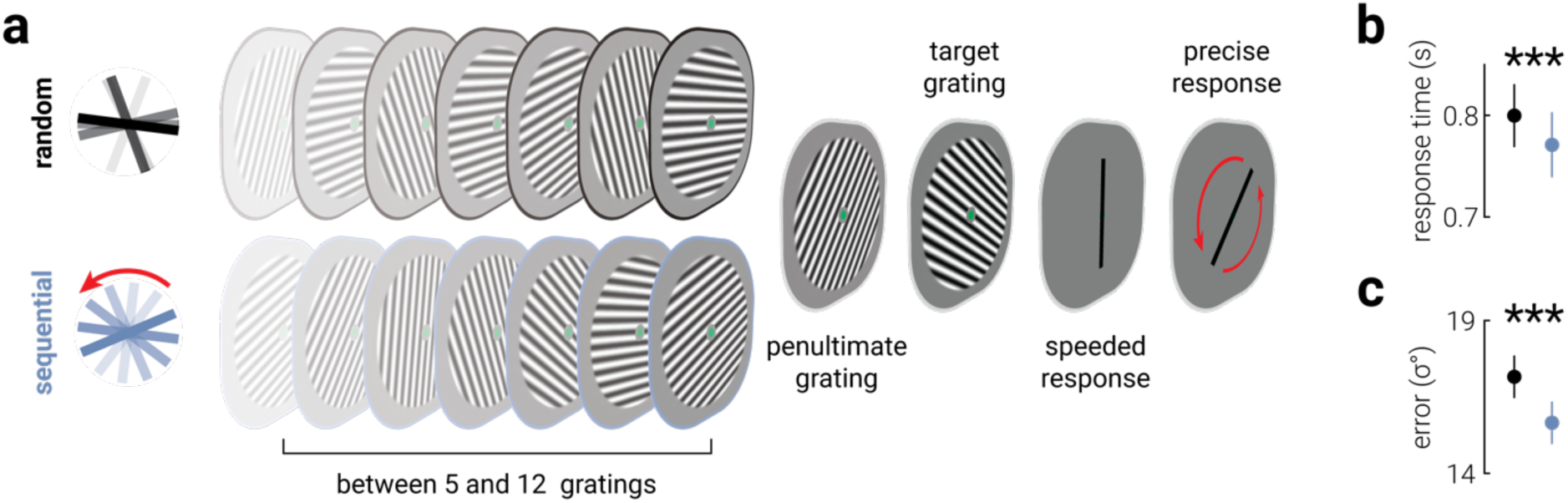
Unexpected stimuli provoke faster and more precise responses. **a**) An illustration of the trial design. Observers viewed a sequence of gratings and were tasked with making a speeded response (clockwise or counterclockwise from vertical) about the target grating before precisely reproducing its orientation. Critically, while the orientation of the target grating was selected at random, the orientations of the preceding gratings were either sequential (rotating at 30° per grating; blue sequence) or random (black sequence). Thus, in the sequential condition, an expectation was established and subsequently violated by the target grating, whereas in the random condition there was no expectation. Note, the changing contrast in the image sequence illustrates the passage of time, and did not occur in the actual stimulus displays. **b, c**) The average (**b**) (speeded) response times and (**c**) (reproduction) precision in the random (black) and sequential (blue) conditions. Asterisks indicate significant differences of *P*<.001.

## METHODS

### Transparency and openness

The data and analysis code for this study are publicly available at https://osf.io/gj7ut/.

### Participants

Sixty self-reported neurotypical human adults (mean±standard deviation age, 24.8±3.9 years; 43 females) participated in Experiment 1 and ten in Experiment 2 (mean±standard deviation age, 24.1±2.3 years; 8 females). Sample sizes were determined by pilot testing and similar experiments within the laboratory. Observers were recruited from The University of Queensland, had normal or corrected-to-normal vision (assessed using a standard Snellen eye chart). All but three participants (authors PD, LJ and ZB) were naïve to the aims of the experiment and all gave informed written consent. The experiment was approved by The University of Queensland Human Research Ethics Committee.

### Apparatus

The experiment was conducted in a dark room. The stimuli were presented on a 32-inch Display++ LCD monitor (Cambridge Research Systems Ltd., Rochester, UK) with 1920 x 1080 resolution and a refresh rate of 120 Hz. Viewing distance was maintained at 48 cm using a chin and head rest, meaning the screen subtended 62.21° x 34.99° (each pixel 3.2’ x 3.2’). Stimuli were generated in MATLAB v2020a (The MathWorks, Inc., Matick, MA) using Psychophysics Toolbox (Brainard, 1997; Kleiner et al., 2007; Pelli, 1997) v3.0.18.13 (see http://psychtoolbox.org/). Gaze direction and pupil diameter were recorded monocularly (right eye) at 1 kHz using an EyeLink 1000 (SR Research Ltd., Ontario, Canada).

### Stimuli, task, and procedure

The stimuli comprised sinewave gratings (1 cycle/°, full contrast, phase randomized between trials) presented centrally within a circular aperture (radius 4°), which was smoothed at the edges, on a mid-grey background. A centrally positioned green fixation dot (radius 0.25°) was presented to reduce eye movements.

As illustrated in **Figure 1a**, trials consisted of 5-12 (counterbalanced across trials) sequentially presented grating stimuli (initial sequence). A variable number of gratings were presented in each trial to motivate observers to attend to the entire sequence. The initial sequence was followed by 2 additional gratings (penultimate and target). The orientations of the gratings in the initial sequence were either randomly selected between 0-180° (random condition) or rotated consistently by ±30° from a random start orientation (sequential condition; rotation direction counterbalanced across trials). The orientations of the penultimate and target gratings were matched between trials of the two conditions, such that for each sequential condition trial there was a random condition trial with the same penultimate and target gratings. In the sequential condition, the penultimate grating orientation was consistent with the rotation of the preceding expectation sequence (i.e., ±30° from the final grating in the sequence), while the target grating was selected at random (1-180°). Each grating was presented for 250 ms, separated by a blank 32 ms inter-stimulus-interval. Following the presentation of the target grating and a 32 ms blank period, a black vertical bar (length 3.2°) was displayed and observers were given 2 s to indicate whether the target grating was oriented clockwise or counterclockwise from vertical using the right or left mouse button, respectively. Upon responding, the black bar was displayed at a random orientation (between 0-180°) and the observers were then required to use the mouse to reproduce the orientation of the target grating (no time limit). Observers were instructed to respond quickly for the two-alternative forced choice (2AFC) task and precisely for the reproduction task. There were 24 trials per block and 16 blocks in total (∼60 min), with equal numbers of random and sequential orientation trials interleaved pseudo-randomly across the session. Participants conducted one to two blocks of familiarization trials, in which feedback was provided on the task, at the start of the session. No feedback was provided in the experiment proper.

Experiment 2 was similar to Experiment 1, except that the spatial position of the gratings varied such that the stimulus appeared to traverse an imaginary circle (radius 6°, step size 12.85°, direction counterbalanced across trials) in the center of the screen. The fixation dot was centered on the moving stimulus, motivating participants to follow the stimulus as it moved around the screen. On half of the trials, the target grating was presented on a cardinal axis aligned to the center of the screen (i.e., above, below, left, or right). The target axis was randomized between blocks and indicated throughout the experiment by a white dot (radius 0.25°) positioned at 10° from center along the corresponding axis; black dots were presented at the same eccentricity on the three remaining axes. On the other half of the trials, the target grating was presented before it reached the target axis. Thus, observers were aware that gratings that were presented on the target axis were target gratings. We opted for this design, rather than a (simpler) temporal cue such as a warning symbol preceding the final grating, to avoid presenting additional unexpected events. The response bar was presented at the location of the target grating. This was repeated 24 times per block and observers performed between 48 and 60 blocks of trials (∼240 min, spread across multiple sessions).

### Eye tracking

Pupil size recordings were epoched to between -500 ms to 1500 ms around target grating onset, normalized to each observer’s average pupil size across the experiment, and a linear model fit to the 500 ms period prior to stimulus presentation was used to baseline each epoch. Missing time samples (e.g., due to blinks) were interpolated. Pupil data from two observers were not collected due to hardware issues and observers with an average pupil diameter >3 standard deviations from the group mean were omitted from further analyses (n=3).

Using the cluster correction analysis described in the *Statistical analyses* subsection below, we found a single period when observers’ pupil diameter was significantly influenced by the experimental manipulation of expectation. To test the relationship between pupil diameter and behaviour, we calculated the average difference in pupil diameter during this period, for each observer, and correlated this with behavioural performance.

### Statistical analyses

Statistical analyses were performed in MATLAB v2020a using CircStat Toolbox v1.12.0.0 (Berens, 2009). Observers’ proportion correct and average response time on the 2AFC (speeded) task was calculated; data from observers who scored <.75 correct across all trials were omitted from further analyses (Experiment 1, n=8; Experiment 2, n=1). Precision on the reproduction task was inferred from the angular standard deviation between the target and response orientations. We only included trials on which the speeded task was answered correctly (91.1%), but this did not change the pattern of results from when all data were included. Repeated measures analysis of variance (ANOVA) and paired *t*-tests were applied to test for differences between conditions. The robust correlation toolbox (Pernet et al., 2013) was used to compute skipped Pearson correlations (default boxplot rule used to identify outliers) to test the relationship between measurements.

When testing the difference in pupil diameter between conditions across time, a cluster correction was applied to mitigate the inflated risk of false positives associated with conducting multiple tests (Pernet et al., 2013). We began by calculating the *t*-statistic at each time point. Next, we calculated the summed value of these statistics (separately for positive and negative values) within contiguous temporal clusters of significant values. We then simulated the null distribution of the maximum summed cluster values using permutation (*n*=1000) of the condition labels, from which we derived the 95% percentile threshold value. Clusters identified in the data with a summed effect-size value less than the threshold were considered spurious and removed. A one-tailed *t*-test was used to test for increased pupil diameter, as we had (a priori) hypothesised that pupil dilation would increase in “surprising” trials. Two-tailed tests were used for all other comparisons.

## RESULTS

### Experiment 1

As shown in **Figure 1b-c**, observers responded faster and more precisely to unexpected grating stimuli than to random stimuli (response time, *t*_51_=5.68, *P*=6.44e^-7^; precision, *t*_51_=5.13, *P*=4.49e^-6^). We tested the inter-individual relationship between response time and precision and found positive correlations in both random and sequential conditions (random, *r*=.58, *P*=5.93e^-6^; sequential, *r*=.55, *P*=2.36e^-5^; **Fig. 2a, b**). This is likely explained by individual differences in factors that influence general task performance, such as fatigue and motivation. To control for these factors and isolate the behavioural changes associated with processing unexpected stimuli, we tested whether the difference in response times associated with unexpected stimuli was related to the difference in precision. We found these changes were positively correlated (*r*=.47, *P*=3.84e^-4^; **Fig. 2c**), suggesting that improvements in response time and precision associated with unexpected stimuli are supported by a common underlying mechanism.

**Figure 2.**
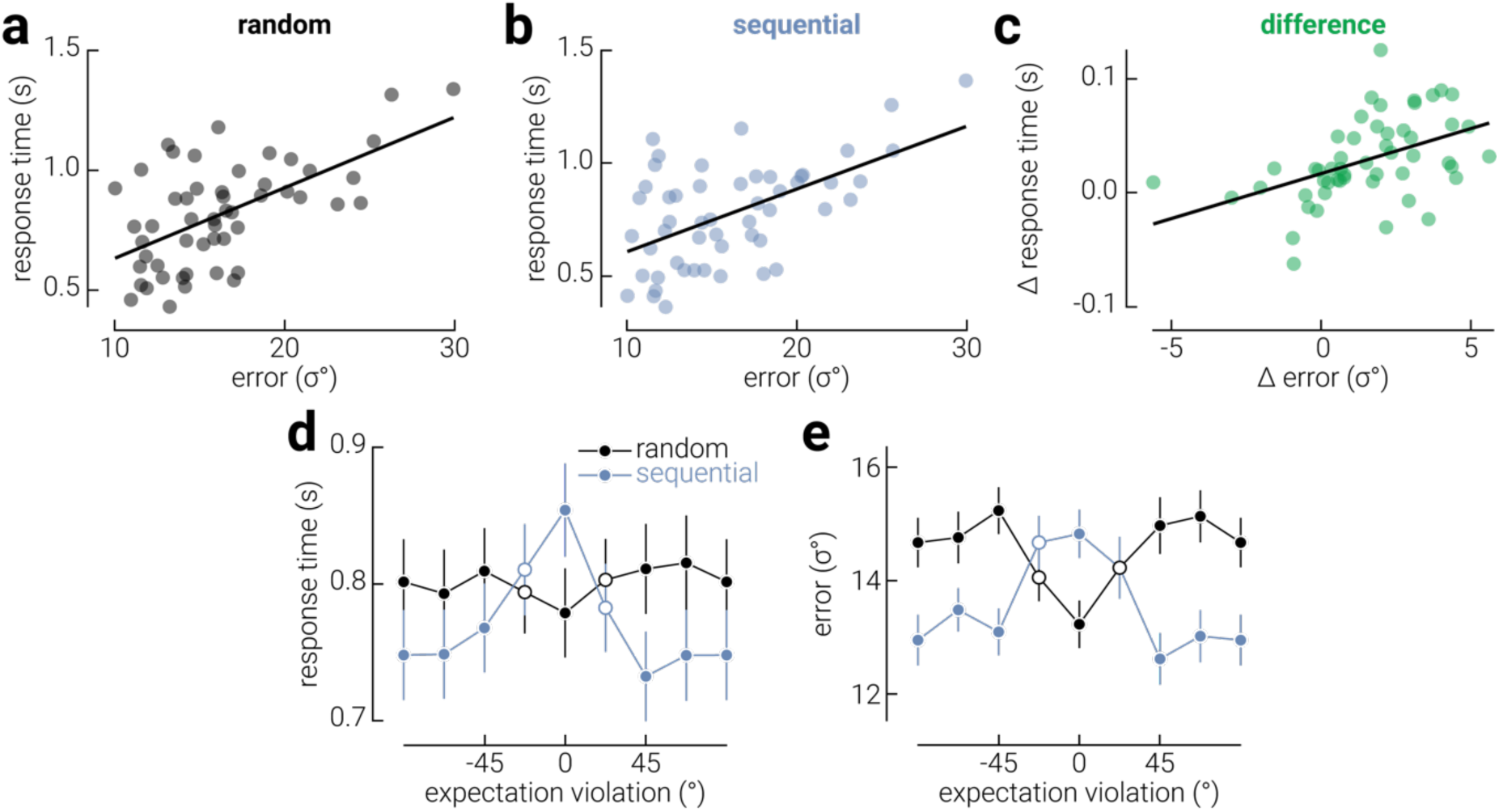
Shared mechanisms and tuning properties for visual predictions. The relationship between response time and precision across observers in the (**a**) random and (**b**) sequential conditions. **c**) The relationship between the difference (random – sequential) in response time and precision across observers. Black lines indicate line-of-best-fit. **d**) Response time and **e**) precision as a function of the magnitude of expectation violation (operationalized as the angular difference between the expected and target orientation) in the random and sequential conditions, binned into eight evenly spaced orientations. Note that while there was no clear expectation in the random condition, the trials in this condition were also grouped according to the angular difference between the penultimate and target grating. In (**d, e**), error bars indicate ±SEM and filled circles indicate magnitudes at which there was a significant difference between conditions.

The orientation of the target grating was randomized, which means that the target violated the expectation established by the preceding gratings on most of the sequential condition trials. However, the magnitude of this violation (operationalized as the angular difference between the expected and target orientation) ranged between ±90°; thus, on some trials the target grating effectively satisfied, rather than violated, this expectation. If improved response time and precision are due to violations of expectation, these improvements should vary as a function of violation magnitude. In line with this reasoning, for both response time and precision, we found a main effect of condition (response time, *F*_1,51_=28.98, *P*=1.86e^-6^; precision, *F*_1,51_=29.74, *P*=1.45e^-6^) and an interaction between violation magnitude and condition (response time, *F*_7,51_=14.66, *P*=1.11e^-16^; precision, *F*_7,51_=14.59, *P*=1.11e^-16^; **Fig. 2d, e**). Pairwise comparisons revealed that response time and precision were worst when expectations were satisfied and best when expectations were violated by >=45°. Indeed, when stimuli were expected (i.e., 0° angular offset in the sequential condition), response time and precision were worse than when there was no established expectation (response time, *t*_51_=5.25, *P*=2.97e^-6^; precision, *t*_51_=4.91, *P*=9.07e^-6^). Thus, the curve formed by the data from the sequential condition effectively represents the tuning function of observers’ predictions. Interestingly, the tuning curve appears to be suppressive, that is, it reflects slower and less precise responses for expected stimuli, consistent with an inhibitory predictive signal. Given that no consistent expectation is generated by random sequences, performance in the random condition should be invariant to the difference between the penultimate and target gratings. However, we observed faster and more precise responses when the penultimate and target gratings were similar. This is likely explained by temporal integration, since the stimuli were presented briefly (250 ms) and in rapid succession (32 ms inter-stimulus interval, ISI). When similar gratings appeared one after another, observers received a clearer sensory signal from which to encode orientation. We note that this finding was incidental to the purpose of our investigation, and responses for unexpected stimuli were nevertheless faster and more precise when compared with the flat profile of responses in the random condition.

To further investigate the reduced precision observed for expected stimuli, we calculated the signed error (bias) as a function of expectation violation (**Supplementary Fig. 1a**). In the sequential condition, there was evidence for a repulsive bias away from the expected orientation. This result is consistent with the reduced precision for expected stimuli. By contrast, we found no clear pattern of bias around the “expected” orientation. We further investigated whether there were differences in task performance between trials in which fewer or more gratings preceded the target in trial, using a median split of the data. For example, stronger predictions may have been established when more spatiotemporally ordered gratings were presented. Indeed, in the sequential condition we found that trials with more preceding gratings were responded to faster and more precisely (**Supplementary Fig. 2**). However, the same result was evident in the random condition, indicating that the effect of sequence length was unrelated to the expectation established by the spatiotemporal order of the stimuli. Rather, this effect appears to be consistent with previous work showing that in tasks with variable stimulus onset asynchrony (SOA), response times are faster for longer SOAs (Niemi & Näätänen, 1981), which is thought to reflect motor priming.

Here and in previous neuroimaging studies of prediction errors (Kok et al., 2012; Tang et al., 2018, 2023), the spatiotemporal structure of visual stimuli was manipulated to establish and violate expectations. It is unclear, however, whether these stimuli sufficiently establish and violate expectation. Pupil dilation is an involuntary physiological marker of arousal/surprise (Preuschoff et al., 2011). If observers’ expectations were violated by the target gratings, we would expect their pupils to dilate in response to these stimuli. For both the sequential and random sequences, pupil diameter increased in response to the target grating, which might have been due to a reduction in luminance between the blank ISI and the target grating, or the target grating and the response bar (**Fig. 3a, b**). Critically, when we examined the difference between random and sequential conditions, pupil diameter was significantly larger following target gratings with larger expectation violations (>±30° from the expected orientation; **Fig. 3c**) from 563 – 1100 ms after target onset. This timing is consistent with previous pupillometry work (Preuschoff et al., 2011), and provides physiological evidence that observers were surprised by gratings that violated the spatiotemporal expectation established in the sequential condition. Critically, there was also a positive relationship between pupil diameter and faster response time in this condition (*r*=.48, *P*=.001; **Fig. 3d**), suggesting a link between violated expectations and improved performance. There was also a positive relationship between pupil diameter and precision but this trend did not reach significance (*r*=.23, *P*=.143; **Fig. 3e**), which may be because there was a longer delay between the unexpected stimulus and the reproduction response than the speeded response. In addition to previous work showing increased cortical activity in response to unexpected events (Den Ouden et al., 2012; Kok et al., 2012; Meyer & Olson, 2011; Todorovic et al., 2011), recent fMRI work has found increased pupil diameter and increased activity in subcortical regions (including the locus coeruleus), which are associated with neuromodulatory systems (dopamine, serotonin, and acetylcholine) (Mazancieux et al., 2023). Our findings complete the characterization of this coordinated response to unexpected events, by revealing the perceptual consequences of potentially related cortical and subcortical activity, and their relationship with physiological markers of surprise.

**Figure 3.**
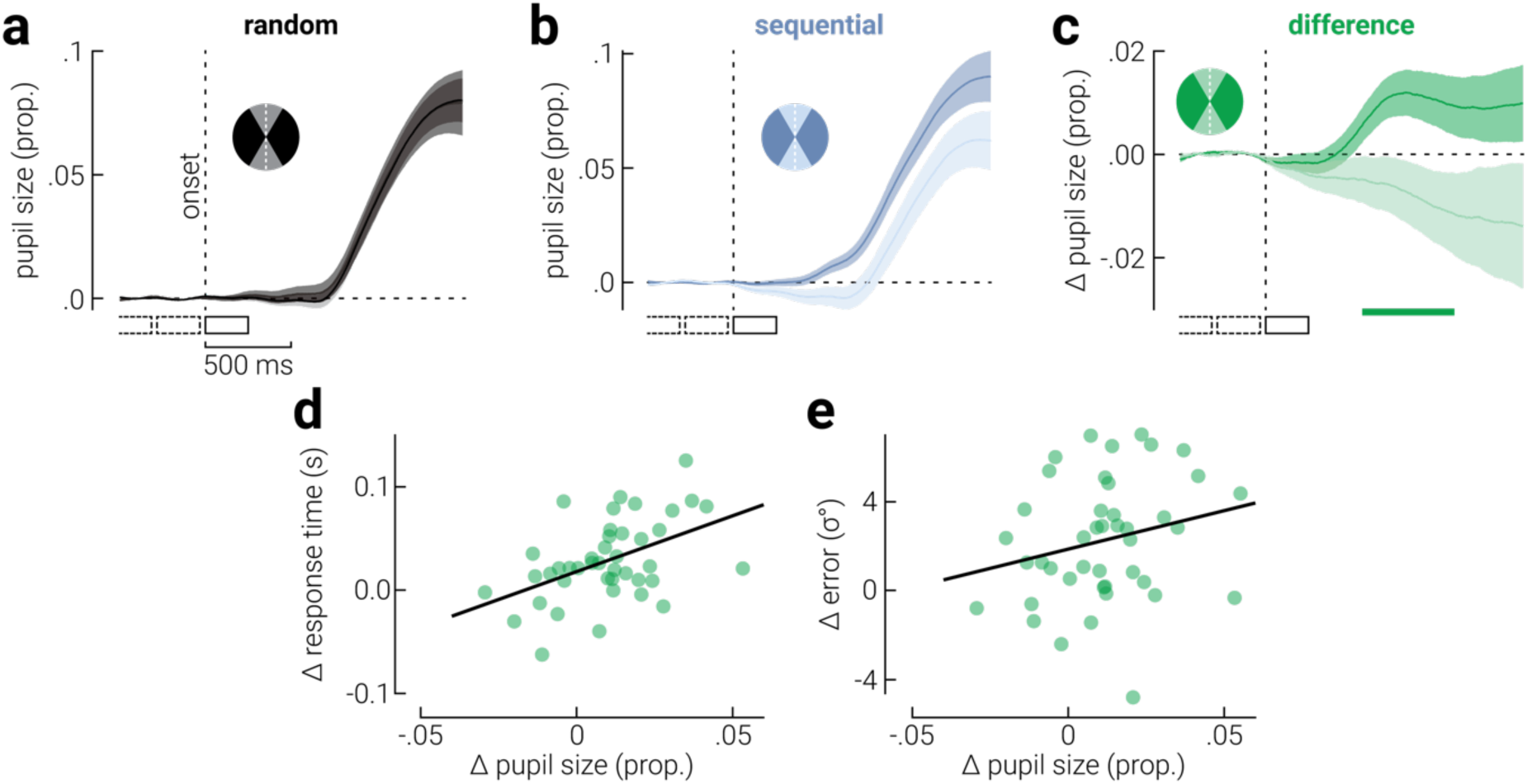
Changes in pupil diameter associated with expectation violation. **a**) Change in pupil size in response to the target grating in the random condition, for trials with smaller and larger than ±30° expectation violation. The inset disk indicates the colours used to show the data for target gratings with orientations that were near to (gray) or far from (black) the expected orientation (white dashed line). **b**) Same as (**a**), but for the sequential condition. **c**) The difference in pupil size between random and sequential condition trials (sequential – random), with larger and smaller expectation violations; colour mapping is the same as in (**a**, **b**). **d, e**) The difference in average response time (**d**) and precision (**e**) between random and sequential conditions as a function of the difference in pupil size. Note that while there was no clear expectation in the random condition, the trials in this condition were also grouped according to the angular difference between the penultimate and target grating. Shaded regions indicate ±SEM and the horizontal bar in (**c**) indicates a cluster corrected period of significant differences between random and sequential conditions. Solid and dashed rectangles in (**a**-**c**) indicate the target and preceding stimulus presentations, respectively.

### Experiment 2

Prioritization of unexpected events may be implemented through neuromodulator-mediated engagement of attention systems (Feldman & Friston, 2010; Hohwy, 2012; Sara & Bouret, 2012). To motivate observers to attend to all presented stimuli, in Experiment 1 we displayed a variable number (5-12) of gratings, such that the identity of the target grating was unknown until the response bar appeared. Thus, on most trials in the sequential condition, both the timing and feature-specific (orientation) properties of the target grating were unexpected. However, in this condition, on trials where the target orientation was noticeably different from the expected orientation, this may have cued participants to the identity of the target. By contrast, when the target orientation was similar to the expected orientation, the identity of the target may have only become apparent upon presentation of the response bar.

To isolate the contribution of the feature-specific violation, we ran a further study (Experiment 2) in which the gratings not only rotated but also traversed an imaginary circle, such that the target grating was either presented at a validly or invalidly cued location (**Fig. 4a**). Thus, on half the trials observers were validly cued to the timing and location of the target grating before it was presented. The target was presented at an invalidly cued location on half the trials to motivate participants to attend to all of the gratings in the sequence, and not just those at the expected location. The invalidly cued condition also served as a replication of Experiment 1, where the timing of the target was also uncertain. Although the cue was only valid on half the trials, participants were aware that the target would never be presented after the cued location, meaning the cue was effectively 100% valid once the grating approached the cued location. We assessed the influence of cueing by comparing performance between random and sequential sequences on trials where the target identity was cued validly or invalidly (**Fig. 4b**).

**Figure 4.**
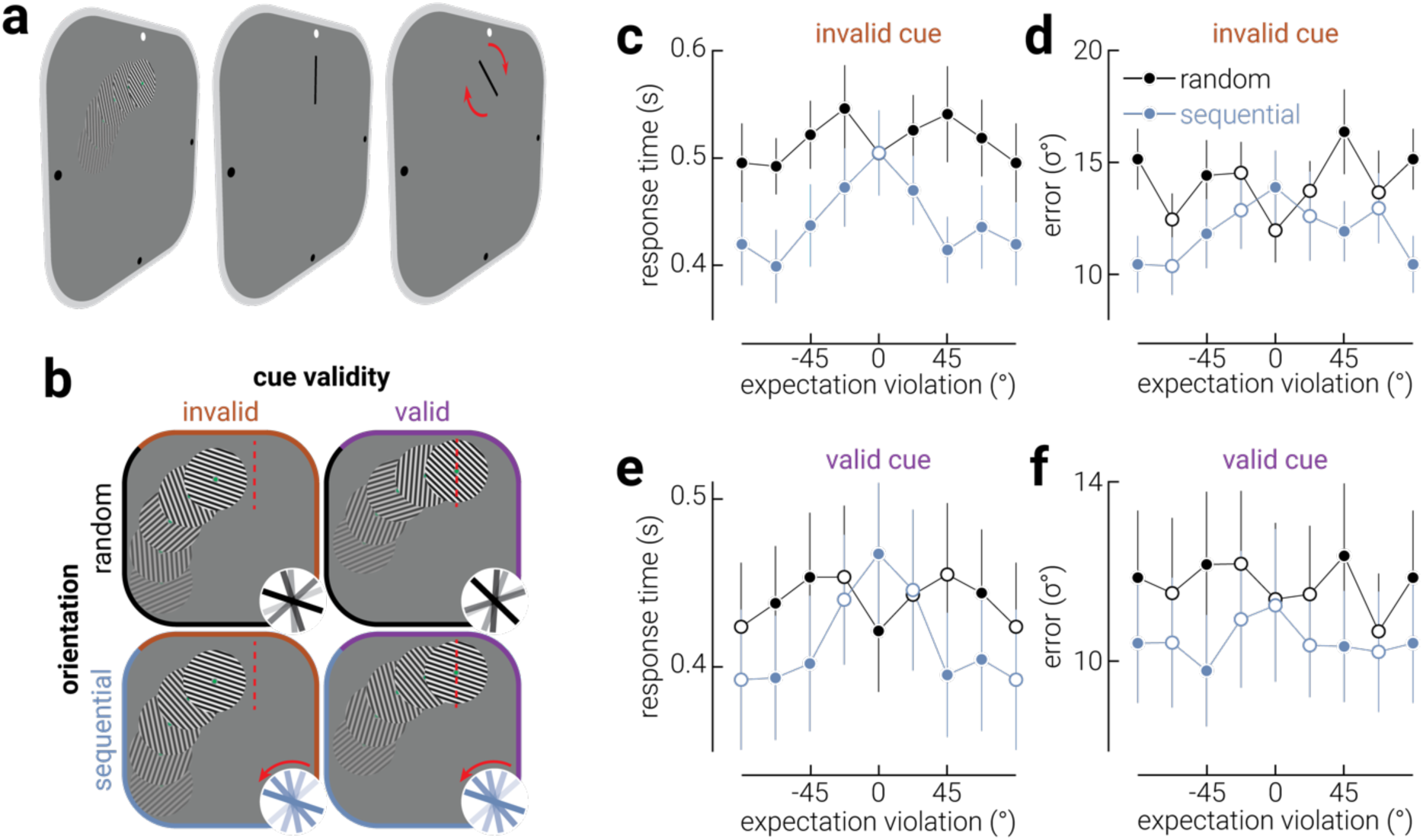
Unexpected stimulus features provoke faster and more precise responses. An illustration of the trial design for Experiment 2, which was the same as in the first experiment, except that stimuli moved around the display such that the target grating was presented either **b**) before (invalid) or at a predefined location (valid, illustrated by dashed red line). Thus, observers were cued to the identity of the target grating before it was presented in the predictable condition. **c**) Response time and **d**) precision as a function of expectation violation, for random and sequential sequences, when the target location was invalidly cued. **e,f**) Same as (**c,d**), but when the target location was validly cued.

For both response time and precision, we found significant effects of target sequence, cue validity, and their interaction (all *P*<.05). Paired *t*-tests of the aggregate data (including all violation magnitudes) revealed that the results from the invalidly cued condition replicated those from the main experiment (response time, *t*_8_=6.91, *P*=1.23e^-4^; precision, *t*_8_=6.88, *P*=1.27e^-4^; **Fig. 4c, d**). Critically, we also found faster and more precise responses in the sequential condition when the target identity was cued validly (response time, *t*_8_=6.06, *P*=3.04e^-4^; precision, *t*_8_=4.84, *P*=.001; **Fig. 4e, f**). Taken together, these results confirm that unexpected features of stimuli (i.e., orientation) are sufficient to produce faster and more precise responses, even when the location and timing of the relevant stimuli are expected.

Given the increased complexity of the task in Experiment 2, relative to Experiment 1 (specifically, the introduction of positional dynamics), one might have expected task performance to be poorer in Experiment 2. For example, participants were required to make regular saccades during the task to follow the location of the stimulus, which might have disrupted visual processing. Despite this, inspection of the data suggests that response times were faster than those in Experiment 2, and reproduction responses were similarly precise. This might be due to the ratio of sample size to number of trials between experiments. In Experiment 1, we collected data from a relatively large sample (n=60), and each participant performed the task for ∼1.5 hours. By comparison, in Experiment 2, we used a smaller number of participants (n=10), and each performed the task for ∼4 hours. Thus, while the task in Experiment 2 may have been more difficult, participants had more opportunity to improve their performance.

## DISCUSSION

Our findings reveal the perceptual consequences of prediction errors and indicate that unexpected events in the environment are prioritized by the visual system both in terms of processing speed and representational fidelity. Prioritizing unexpected events likely supports immediate and future adaptive behaviours through multiple operations, such as rapid responses to potentially harmful events and precise updating of internal predictive models (Friston, 2009; Soltani & Izquierdo, 2019), and may be operationalized through subcortically regulated attention systems (Feldman & Friston, 2010; Hohwy, 2012; Sara & Bouret, 2012).

The finding that unexpected events are reproduced more precisely is consistent with EEG data from humans (Tang et al., 2018) and Ca^2+^ imaging data from mice (Tang et al., 2023), but appears to contradict some previous fMRI data in humans (Kok et al., 2012). These discrepancies could be due to biophysical differences between the imaging signals used in these studies. Specifically, blood-oxygen-level-dependent (BOLD) signals may not have the temporal resolution to disambiguate pre-activation (Kok et al., 2017) from prediction error signals. Pre-activation likely biases performance toward expected outcomes and thus may obscure the suppressive effects of prediction errors on these same outcomes. Under this interpretation, our findings could indicate that perception reflects an interaction between prediction and sensory processing input, rather than pre-activation, which may facilitate attentional deployment.

How do our findings fit with previous behavioural studies? Previous work found better perceptual sensitivity for stimuli that complied with spatial and temporal cues (Rohenkohl et al., 2014). However, this may be due to differences in attention prior to presentation of attended and unattended stimuli, rather than the influence of prediction error. That is, cueing attention to a particular location that is either valid or invalid, relative to the spatiotemporal location of the upcoming stimulus, creates an attentional bias between conditions before the expected/unexpected stimulus is presented. By contrast, in our experiments, spatial and temporal attention were equally focused on expected and unexpected stimuli (at least prior to stimulus presentation).

Sequential dependency paradigms, which establish expectations differently from those typically used to study prediction errors, show that expected events are responded to faster and more accurately than unexpected events (Remington, 1969). By contrast, here we found that observers responded faster to unexpected than expected stimuli. This suggests distinct mechanisms drive the behavioural consequences of expectancy associated with classic sequential dependency paradigms and those observed here. Indeed, our finding that unexpected events are processed faster provides some support for the claim that the differences observed in sequential dependency paradigms are a result of motor priming, rather than a change in sensory encoding (Jentzsch & Sommer, 2002). Abreo et al. (2023) used a similar design to that employed here (**Supplementary Fig. 3**), but concluded that expected gratings were reproduced more precisely, which seems to be in direct conflict with our findings. However, re-analysis of their data revealed that their results may have been driven by temporal integration of the penultimate and target gratings, and that the number of stimuli used by Abreo et al. (4-7) – approximately half the number used here (6-13) – did not establish a sufficiently reliable expectation.

In summary, here we resolve inconsistent findings on the influence of prediction errors on sensory encoding by showing that unexpected events are responded to faster and reproduced more precisely than those that are either expected or occur randomly and thus without any explicit expectation. These findings offer empirical evidence for neural prioritization of unexpected events, which support immediate adaptive responses and updating of future predictions.

## Acknowledgements

This work was supported by an Australian Research Council (ARC) Discovery Early Career Researcher Award awarded to RR (DE210100790). RR was also supported by a National Health and Medical Research Council (NHMRC; Australia) Investigator Grant (2026318). JBM was supported by an NHMRC Ideas Grant (1165337) and an NHMRC Investigator Grant (2010141).

## Footnotes

1 Representational fidelity refers to the accuracy of the memory trace of the target orientation. Higher fidelity is indicated by lower errors in a reproduction task.

