## Supplementary material for "Violated predictions enhance the representational fidelity of visual features in perception"

To further investigate the reduced precision observed for expected stimuli, we calculated the signed error (bias) as a function of expectation violation (**Supplementary Fig. 1a**). We then used a bootstrap approach to quantify serial dependence in sequential and random conditions (Abreo et al., 2023). For both conditions, data were fit with a von Mises derivative function with a mean of zero;

$$f(x) = a\Delta\phi_{\sigma} + c$$

where $a$ denotes amplitude, $\phi_{\sigma}$ denotes the Von Mises distribution with mean of zero and kappa $\sigma$, and $c$ is a constant offset. This function was used, rather than a difference of Gaussian function (Fischer & Whitney, 2014), because the data were sampled across the full range of values within a circular coordinate space. We fit the von Mises function to group-averaged data using a random subsample (with replacement) of observers on 20,000 bootstrap permutations. For each permutation, the amplitude and offset parameters of the fit were saved, generating a distribution of parameters which were converted to *P* values by dividing the number of bootstrap samples above or below zero by the number of permutations (20,000) and multiplying by two (two-tailed, nonparametric test of the amplitude parameter against zero). In the sequential condition, two patterns of bias were qualitatively apparent in the results. First, responses were not distributed evenly around zero; rather, there was an overall bias toward negative errors (sequential, $P$=.0286; random, $P$=.0006). This suggests that participants exhibited a bias in the opposite direction to the rotation of gratings, and appears to be independent of expectation. The second pattern was a repulsive bias away from the expected orientation ($P$=.0292). This result is consistent with suppression of the expected orientation, and inconsistent with a (Bayesian) integration account, which would predict better precision coupled with an attractive bias around the expected orientation. By contrast, while the same overall negative bias was apparent in the random condition, there was no clear pattern of bias around the "expected" orientation ($P$=.5120). Related to these results, previous work has shown that responses on reproduction tasks can be biased toward previously presented items, a phenomenon referred to as *serial dependence* (Cicchini et al., 2014; Fischer & Whitney, 2014). To test whether serial dependence influenced responses in our experiment, we calculated bias as a function of the angular distance between the penultimate and target gratings (**Supplementary Fig. 1b**). Note that this serial dependence analysis is similar to the analysis of bias around the expected orientation, but it does not account for the direction of rotation of the preceding stimuli in the sequence. In the sequential condition, we observed marginal evidence for serial dependence, i.e., attraction toward the orientation of the penultimate grating (*P*=.0682, one-tailed test). Existing serial dependence literature has shown that this attractive bias is coupled with increased precision at small offsets between penultimate and target items (Cicchini et al., 2024). By contrast, we found a repulsive bias away from the expected orientation in the sequential condition, which is consistent with reduced precision for expected items. Thus, our measurements may underestimate the influence expectancy on perception, as the influence of expectation on task performance appears to directly conflict with serial dependence in our design.

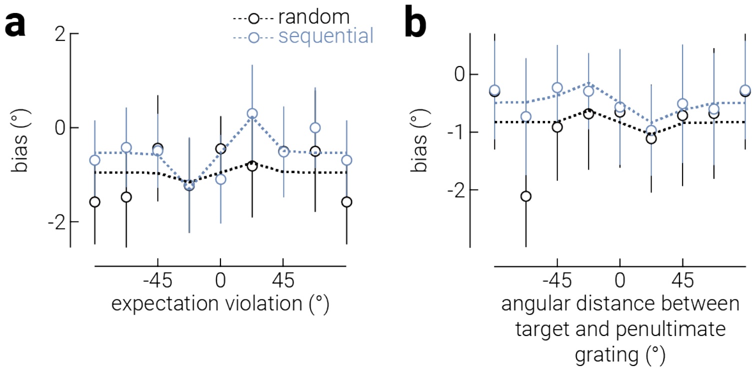


**Supplementary Figure 1. Reproduction biases. a**) Signed error (bias) of reproduction responses as a function of expectation violation in Experiment 1. **b**) Same as (**a**), but as a function of the angular distance between the penultimate and target grating. Dashed lines indicate derivative Von Mises functions fit to the average data. Error bars indicate ±95% confidence intervals. We note there is a large negative bias in the random condition at -57.5°, which we think is likely due to chance, as it is not present in the sequential condition, nor is it accompanied by unusual response times or precision at this offset.

In Experiment 1 we used a variable number of preceding gratings in the trial sequences, in part to motivate participants to attend to all gratings in the possibility that they might be a target. It is possible that performance on the task varied as a function of the number of gratings in the sequence. For example, stronger predictions may have been established when more spatiotemporally ordered gratings were presented. To investigate how the number of gratings in the sequence influenced task performance, we split the data from the sequential condition of Experiment 1 according to whether: (a) the target orientation was “expected”, i.e., within ±30 ° of the expected orientation, or “unexpected”, i.e., more than 30° from the expected orientation, and (b) there were eight or fewer presentations versus nine or more presentations in the preceding sequence (median split). We then compared the response time and error between these groups. The results of this analysis are shown **Supplementary Figure 2**.

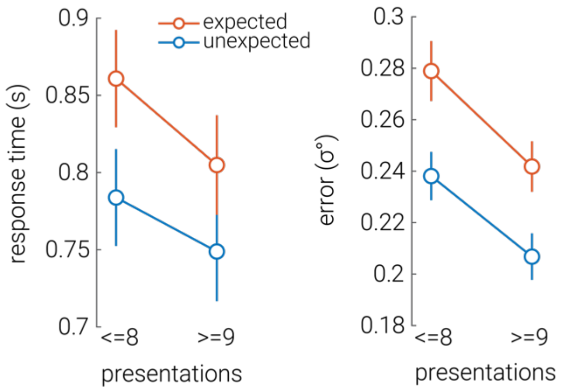


**Supplementary Figure 2.** Response time and reproduction error for “expected”, i.e., within ±30° of the expected orientation, or “unexpected”, i.e., more than 30° from the expected orientation, targets in the sequential condition of Experiment 1, for trials in which there were fewer (<=8) or more (>=9) presentations in the sequence.

Consistent with our main analyses, we found that unexpected targets were responded to faster and more precisely (response time, $F_{1,51}$=76.93, P=9.25e-12; precision, $F_{1,51}$=70.90, P=3.27e-11). We also found that targets on trials with more preceding gratings were responded to faster and more precisely (response time, $F_{1,51}$=28.99, P=1.86e-6; precision, $F_{1,51}$=26.93, P=3.68e-6). However, we found no evidence of an interaction between these effects (response time, $F_{1,51}$=2.61, P=.112; precision, $F_{1,51}$=0.29, P=.593). The finding that response times were faster on trials with more presentations is consistent with previous work showing that in tasks with variable stimulus onset asynchrony (SOA), response times are faster for longer SOAs (Niemi & Näätänen, 1981), which is thought to be an outcome of motor priming. The finding that precision was better on trials in which there were more presentations may reflect increased attention to stimuli that were presented later in the sequence (Woodrow, 1914).

Abreo et al. (2023) used a similar design to that employed in Experiment 1 of the current study (**Supplementary Fig. 3a**), but concluded that expected gratings were reproduced more precisely than random or unexpected gratings, which seems to be in direct conflict with our findings. How can we resolve this discrepancy? To understand the discrepancy between our results and those of Abreo et al. (2023), we re-analyzed their data. We computed response precision as a function of expectation violation magnitude (unexpected condition) and angular distance between penultimate and target grating (random condition). We also computed precision in the expected condition, which is the same as data in the unexpected condition, at zero expectation violation.

As shown in **Supplementary Figure 3b**, while there was a significant effect of violation magnitude ($F_{1,35}$=15.96, $P$=3.61e$^{-4}$), there was no effect of (random/unexpected) condition ($F_{7,35}$=0.78, $P$=.384) and a borderline significant interaction ($F_{7,35}$=2.05, $P$=.049). These results may indicate that the number of stimuli used by Abreo et al.) (4-7) – approximately half the number used here (6-13) – did not establish a sufficiently reliable expectation. This is further supported by the precision in the expected condition of their study (**Supplementary Fig. 3b**, pink marker), which is equivalent to the precision in the matched violation magnitude category for the random and unexpected conditions. Rather, the difference in precision across violation magnitudes in the study of Abreo et al. (2023) is likely the result of temporal integration of the penultimate and target gratings. This phenomenon appears to be less prominent in the random condition of our experiment, probably because Abreo et al. (2023) backward masked their target stimulus. Backward masking reduces the time afforded to process the target, thus increasing the relative benefit of congruent temporal integration with the penultimate grating. This backward masking also likely explains why the responses of their observers were less precise, on average, than those in our study.

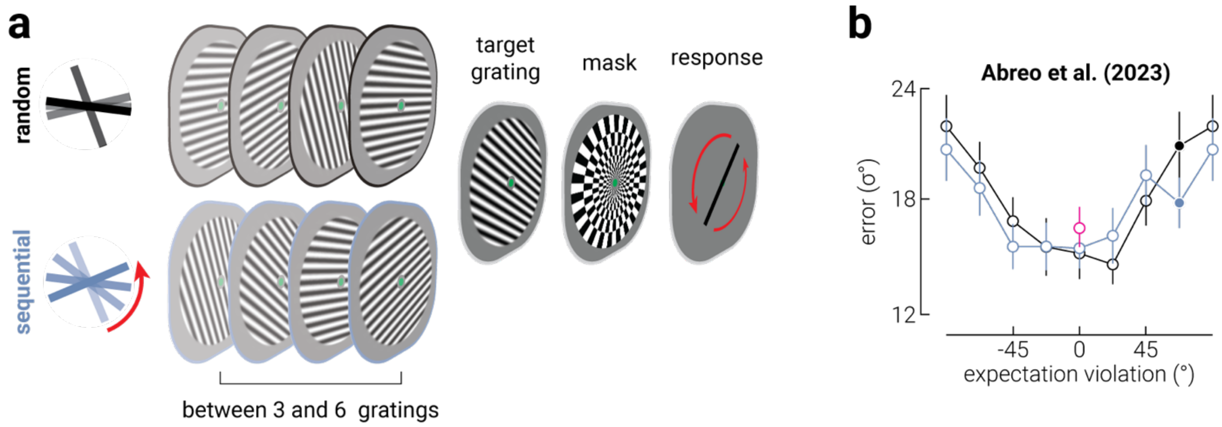


**Supplementary Figure 3. Re-analysis of published data supports prioritization of unexpected events. a**) An illustration of the trial design used by Abreo et al. (2023), which was the same as that used in our first experiment, except they: 1) used fewer gratings to establish an expectation, 2) backward masked the target grating, and 3) did not include a speeded response. **b**) Re-analysis of data from Abreo et al. (2023) showing precision as a function of the magnitude of expectation violation (operationalized as the angular difference between the expected and target orientation) in the random and sequential conditions. The pink marker indicates data from a third "expectation" condition, where the target grating always satisfied the expectation established in the preceding sequence. Note that while there was no clear expectation in the random condition, the trials in this condition were also grouped according to the angular difference between the penultimate and target grating. Filled circles indicate magnitudes at which there was a significant difference between conditions.
